# Multimodal social context modulates behavior in larval *Drosophila*

**DOI:** 10.1101/2025.03.24.644986

**Authors:** Akhila Mudunuri, Élyse Zadigue-Dubé, Katrin Vogt

**Affiliations:** Department of Biology, University of Konstanz, Konstanz, Germany; Centre for the Advanced Study of Collective Behaviour, University of Konstanz, Konstanz, Germany; Max Planck Institute of Animal behavior, Konstanz, Germany

**Author notes:** Department of Biology, McGill University, Montréal, Quebec, Canada (Élyse Zadigue-Dubé).

## Abstract

All animals need to navigate and make decisions in social environments. They influence each other’s behavior, but how important this is and how they process and represent social information in their brain is less well understood. This includes fruit flies and fly larvae, which are usually not known as “social insects”. Using a *Drosophila* larva assay with reduced stimulation, we found that larval groups show enhanced dispersal and distance from each other in the absence of food. This social context-dependent modulation overrides responses to other external sensory cues and is shaped by developmental social experience. Leveraging the genetic toolbox available in *Drosophila*, we find that different sensory modalities are required for normal social context modulation. Our results show that even less social animals like fly larvae are affected by conspecifics and that they recognize each other through multimodal sensory cues. This study provides a tractable system for future dissection of the neural circuit mechanisms underlying social interactions.

## Introduction

Navigating social interactions is essential in any individual’s life and can have strong implications for fitness and survival. Living with conspecifics can be advantageous as it allows for shared and better access to environmental information, facilitating access to resources (*1*, *2*) or improving predator avoidance (*3*, *4*). However, group living also increases competition for mates and limited food sources (*5*, *6*). Therefore, decision-making in a social context, in interactions with other animals and conspecifics, is complex and requires the integration of all available information from the environment. Social context is not only omnipresent and critical for decision-making, but it is also required for healthy development (*7*). As the brain evolved in a social world, further investigation into the social neural circuits is necessary to understand its full functionality.

Understanding how conspecifics sense each other is a good starting point for understanding social behavior from a neuroscience perspective. In locusts, mechanosensory interactions play a crucial role in swarm formation (*8*, *9*), mosquitoes integrate visual and acoustic information to swarm (*10*) and ants rely on both mechanosensation and chemosensation to communicate (*11*). Zebrafish primarily seem to use vision to interact with conspecifics (*12*), whereas pheromones trigger sexual or aggressive behaviors in mammals (*13*). Despite these insights, we know little about the underlying neural circuitry that processes this sensory information. Studying social behavior in *Drosophila* is promising, as the available genetic toolkit and the whole-brain connectome make it a tractable system for dissecting the circuit mechanisms underlying conspecific interactions.

*Drosophila melanogaster*, even though not considered a social insect, is exposed to conspecifics across all developmental stages. Flies exhibit local aggregation even in the absence of external stimuli (*14–16*) and aggregate on food to mate and oviposit (*17*, *18*). Similarly, *Drosophila* larvae socially aggregate to dig together into food substrate to reach better resources (*19*). This collective digging occurs during their feeding stage; however, when entering the wandering state to look for a pupation site, aggregation behavior decreases (*20*). Larval aggregation can also be beneficial as they can collectively defend themselves against a harmful fungus (*21*). However, when food is scarce or under extreme crowding, they need to compete for resources and can even resort to cannibalism (*22*). Thus, flies and fly larvae face many complex decisions when interacting with conspecifics and might need to integrate social information.

In flies, multisensory cues mediate social interactions. Mechanosensory neurons are necessary to convey social information to enhance a collective odor avoidance response (*23*). The formation of spontaneous fly clusters in the absence of external stimuli requires multiple sensory modalities such as vision, audition, olfaction, taste, and touch (*15*). Flies can learn to become sociable by recognizing conspecific smell and integrating this information within the mushroom body (*24*). How larvae sense each other and how larval-larval interactions are affected by changes in environment or prior social experience is less studied. Larvae require vision to coordinate collective digging in food, to be attracted to moving conspecifics, and to avoid collisions (*19*, *25*, *26*). Other sensory systems also seem to be involved in modulating social interactions. Collective digging requires the mechanosensory channel NompC (No mechanoreceptor potential C) (*19*). Larval deposits induce attraction via the ppk23 (pickpocket 23) receptor (*27*). However, larvae suppress cannibalism toward fly eggs coated with a maternal pheromone, which is also sensed by the ppk23 receptor (*28*). Thus, larval social interactions also seem to be complex, context-dependent, and involve multiple sensory systems.

In this study, using a well-controlled and stimulus-reduced behavioral assay, we show that fly larvae disperse away from conspecifics in the absence of any external stimuli. This robust avoidance behavior is prioritized over responses to other external stimuli, such as aversive light and attractive fructose. During development, larvae seem to get used to social context, as larvae raised in isolation from the egg show increased dispersal from conspecifics. Investigating the sensory systems involved in conspecific avoidance, we show that multiple sensory modalities are required for normal social recognition and avoidance behavior. Our results highlight the importance of social context on behavior and the complexity of social sensory cues provided by conspecifics.

## Results

### *Drosophila* larvae disperse away from conspecifics

To understand larval-larval interactions, we tested groups of 15 Wild-type Canton-S larvae in a square arena (25 cm x 25 cm) containing 2% agarose in the dark for 10 minutes (Movie S1, Fig. 1A). Larvae started in the middle of the arena, their behavior was recorded and analyzed using custom tracking software (*29*). As a control for biases in the arena (border preference) or locomotion defects, we also tested individual larvae and randomly assigned 15 individuals into an artificial superimposed group for further analysis (Movie S2, Fig. 1B). We analyzed the first 100 seconds of the experiment in detail to study their dispersal behavior. Individual larvae tend to make shorter runs and have a higher turn rate compared to the group (Fig. 1B/Fig. S1A). Analysis of the mean squared displacement (MSD) from the center shows that larvae in the group disperse more than individual larvae (Fig. 1C). This influence of the social context in the group persists over the whole time of the experiment: larvae in groups have higher MSD than individual larvae (Fig. S1B). We visualized larval distribution in a heatmap by plotting the difference in probability of finding a grouped larva to an individual larva in a certain arena location (Fig. 1C). Individuals stay in the middle (blue squares). Grouped larvae disperse further away from the center (red squares). This trend also persists over the whole experiment (Fig. S1C).

**Fig. 1:**
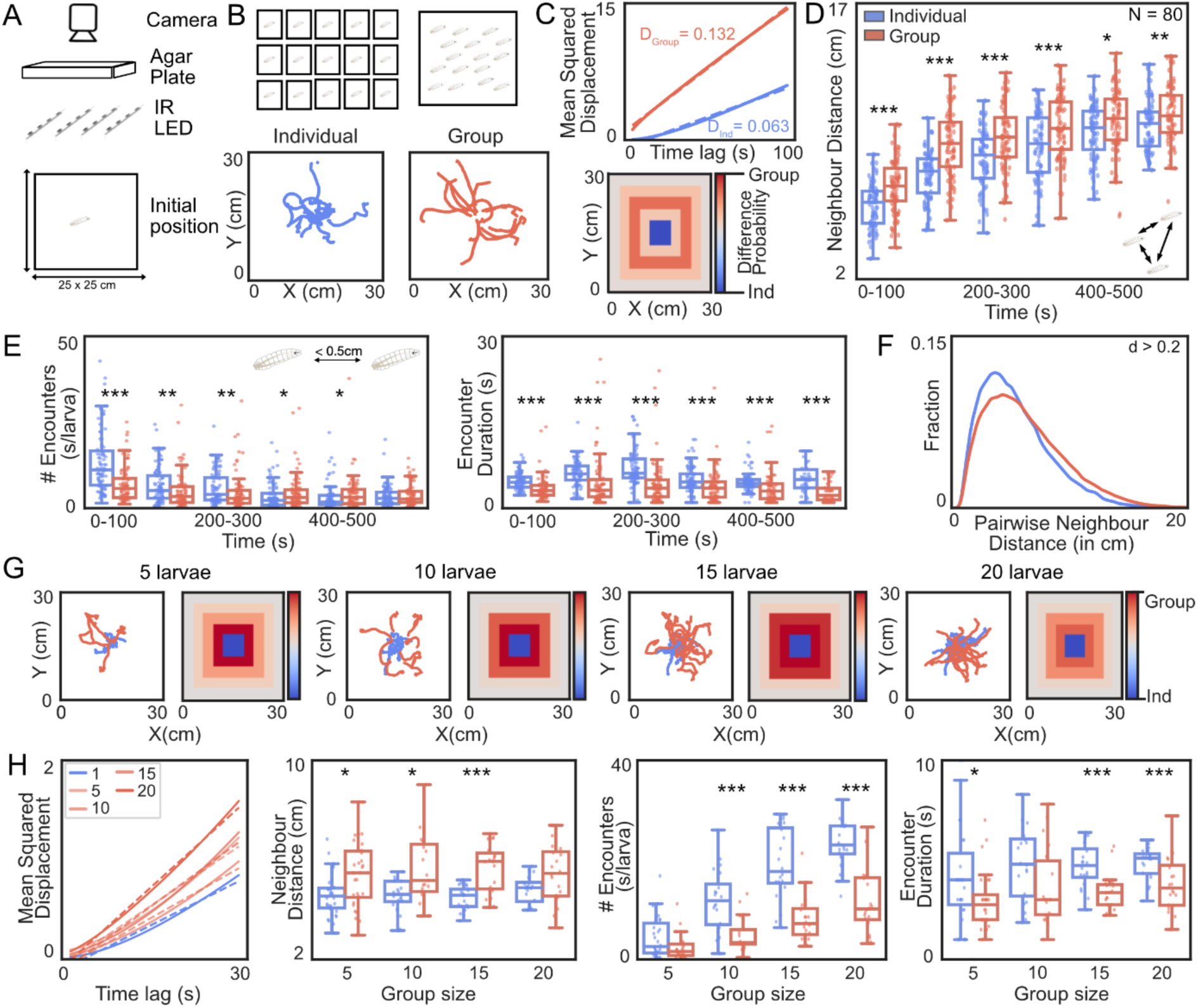
Drosophila larvae in a group show enhanced dispersal: (**A**) Larvae were placed in the center of a square plate (25 x 25cm) containing 2% agar and recorded for 10 min. with infrared illumination. (**B**) Individuals or groups of 15 larvae were tested. Tracks of 15 individuals were superimposed to form a control group. Sample trajectories for the first 100s of an experiment for a superimposed (blue) and a real group (red). (**C**) Mean squared displacement (MSD) for all real and superimposed groups (N = 80 trials per treatment) was plotted as a function of time interval for the whole experiment. The heatmap visualizes the spatial distribution of larvae for the first 100s by subtracting the probability of finding an individual larva from a real group larva. Blue indicates an abundance of individual larvae, and red indicates an abundance of real group larvae. (**D**) The average neighbor distance (ND) between all pairs in a real or artificial group over time is shown. (**E**) Real encounters are less frequent and shorter than artificial encounters in the superimposed group. (**F**) Pairwise neighbor distances for the first 100s in superimposed (blue) and real (red) groups (effect size = Cohen’s d (0.2 < d < 0.5 – small). (**G**) Trajectories and heatmaps of larval groups of 5 (N =30), 10 (N =20), 15 (N =20), and 20 (N = 20) animals, compared to artificial groups with the same sizes. (**H**) MSD and ND of all group sizes for the first 100s. Encounter frequency and duration of real group larvae are lower across all tested group sizes. (Mann-Whitney U test = * p < 0.05; ** p < 0.01, *** p < 0.001).

We asked whether larvae in groups generally disperse in the arena, even when presented with a food context. When adding fly food to the arena center, the group however stays in the food (Movie S3, Fig. S1D). To quantify social interactions between larvae in the empty arena, such as clustering, we analyzed neighbor distance (ND), the average distance between all larval pairs in artificial and real groups of 15 larvae.

Over 10 minutes, the ND of the real group steeply increases and plateaus after a few minutes (Fig. 1D). In the artificial group, the ND slowly and gradually increases over 10 minutes. Overall, the ND of the real group is significantly higher than the ND of the artificial group. This is consistent over the first 200s for most combined wild-type experiments in this dataset (Fig. S1E). To quantify how often and how long larvae interact locally, we counted close encounters with less than 0.5 cm distance between two larvae (Fig. 1E). This encounter frequency is generally lower in the real group compared to the artificial group and larvae in real groups also spend less time in an encounter. During an encounter, larvae in real groups pause and make big turns away from each other, whereas larvae in the artificial group with no real encounters show no change in behavior (Fig. S1F). Larvae start in the middle and thus encounter conspecifics most often in the first 100s (Fig. 1E). Within this time frame, pairwise neighbor distances (PD) are higher in the real group, supporting further that grouped larvae do not cluster but spread out evenly to avoid conspecifics (Fig. 1F). Over the whole experiment, the PDs in the artificial group become more similar to the values of the real group (Fig. S1G).

We hypothesized that larvae might disperse more in the group due to collisions with conspecifics within the first 100s. Adding non-larval obstacles, small agar pieces, to the arena did not increase dispersal in individual larvae but rather attracted them to stay in the arena center (Movie S4, Fig. S1H). Thus, the encounter with a conspecific, and not just any collision, enhances dispersal.

Testing groups of different sizes (5, 10, 15, and 20 larvae), we find the same social context modulation. Individual larvae stay in the center, whereas grouped larvae spread out more (Fig. 1G). The MSD and ND in all groups are higher than in the individuals (Fig. 1H). Also, conspecific encounters in all real groups are less frequent and shorter than in the artificial groups of the same size (Fig. 1H). All groups were tested in the same arena, thus the available space for larvae to spread out is limited in bigger group sizes. We, therefore, chose a group size of 15 for all further experiments.

### Social context modulation overrides responses to external stimuli

Social context modulates the behavior of larvae in an empty experimental arena. As groups rarely interact in an unstimulated environment, we next exposed larvae to two different external stimuli. First, larvae were tested on an evenly distributed 2M fructose/agar substrate, which is usually attractive (*30*) (Fig. 2A). The social modulation was reproducible on all substrates; larvae in the group dispersed more than individual larvae (Fig. 2B). ND and PD suggest that individuals stay closer together on fructose, potentially as they dwell in the center due to fructose attraction; real groups however behave the same on both substrates (Fig 2. C/D).

**Fig. 2:**
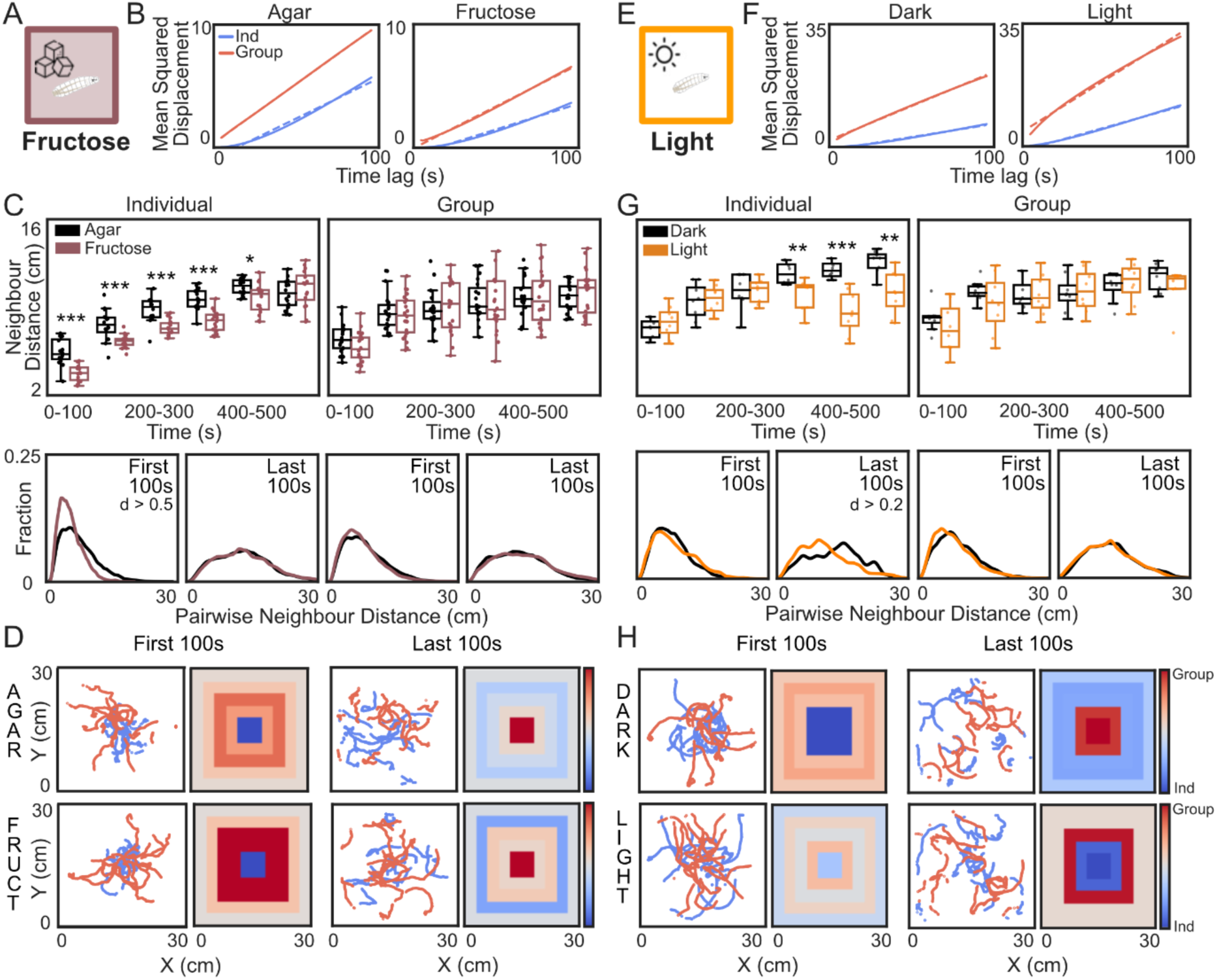
Social context modulation overrides responses to external stimuli: (**A**) Larvae were tested on a 2M Fructose/Agar substrate. (**B**) Mean squared displacement (MSD) for each treatment, comparing superimposed groups (N = 16 per treatment, blue) and real groups (N = 20 per treatment, red). (**C**) Top: Average neighbor distance (ND) over time for the superimposed and real groups. Bottom: Pairwise neighbor distance (PD) for the first and last 100s of the 10-minute experiment. Left: superimposed groups, Right: real groups (Effect size = Cohen’s d ( 0.2 < d < 0.5 – small; 0.5 < d < 0.8 – moderate). black = agar, brown = fructose. (**D**) Sample trajectories and distribution heatmaps (superimposed groups = blue, real groups = red). (**E**) Larvae were tested in an arena with white light LEDs at the arena border. (**F**) MSD for each treatment, comparing superimposed groups (N =8 per treatment, blue) and real groups (N = 8 per treatment, red). (**G**) Top: Average ND over time for the superimposed and real groups. Bottom: PD for the first and last 100s of the 10-minute experiment. Left: superimposed groups, Right: real groups. black = light OFF, orange = light ON (**H**) Sample trajectories and distribution heatmaps (superimposed groups = blue, real groups = red). (Mann-Whitney U test = * p < 0.05; ** p < 0.01, *** p < 0.001).

Second, larvae were tested on a pure agar substrate illuminated with white light LEDs from the arena borders, a stimulus they usually avoid (*31*) (Fig. 2E). The social context modulation was reproducible in the light and dark arena; larvae in the group dispersed more than individual larvae (Fig. 2F). In the first half of the light-on experiment, individual larvae showed high dispersal potentially due to light exposure arousal (Fig. 2G/H). In the second half of the experiment, they even seemed to avoid the bright borders with the LEDs and moved toward the arena center, resulting in a low ND. However, no such light avoidance response was visible in the real group (Fig. 2G/H).

### Developmental experience influences social modulation

We next asked whether social context modulation is an innate behavioral response or whether it is affected by prior conspecific exposure. Therefore, we reared larvae in isolated or highly crowded conditions and compared those to standard conditions (Fig. 3A). For isolation, we placed individual eggs in separate wells (48-well plates) filled with fly food, allowing them to develop without any conspecific interaction throughout development. For highly crowded conditions, we placed 50 fly eggs in a single well (48-well plate) filled with fly food. This condition created a competitive environment due to limited food availability, resulting in a mortality rate of approximately 50%. MSD revealed that isolated larvae disperse more when tested in groups than tested individually, whereas larvae from crowded conditions disperse less (Fig. 3B).

**Fig. 3:**
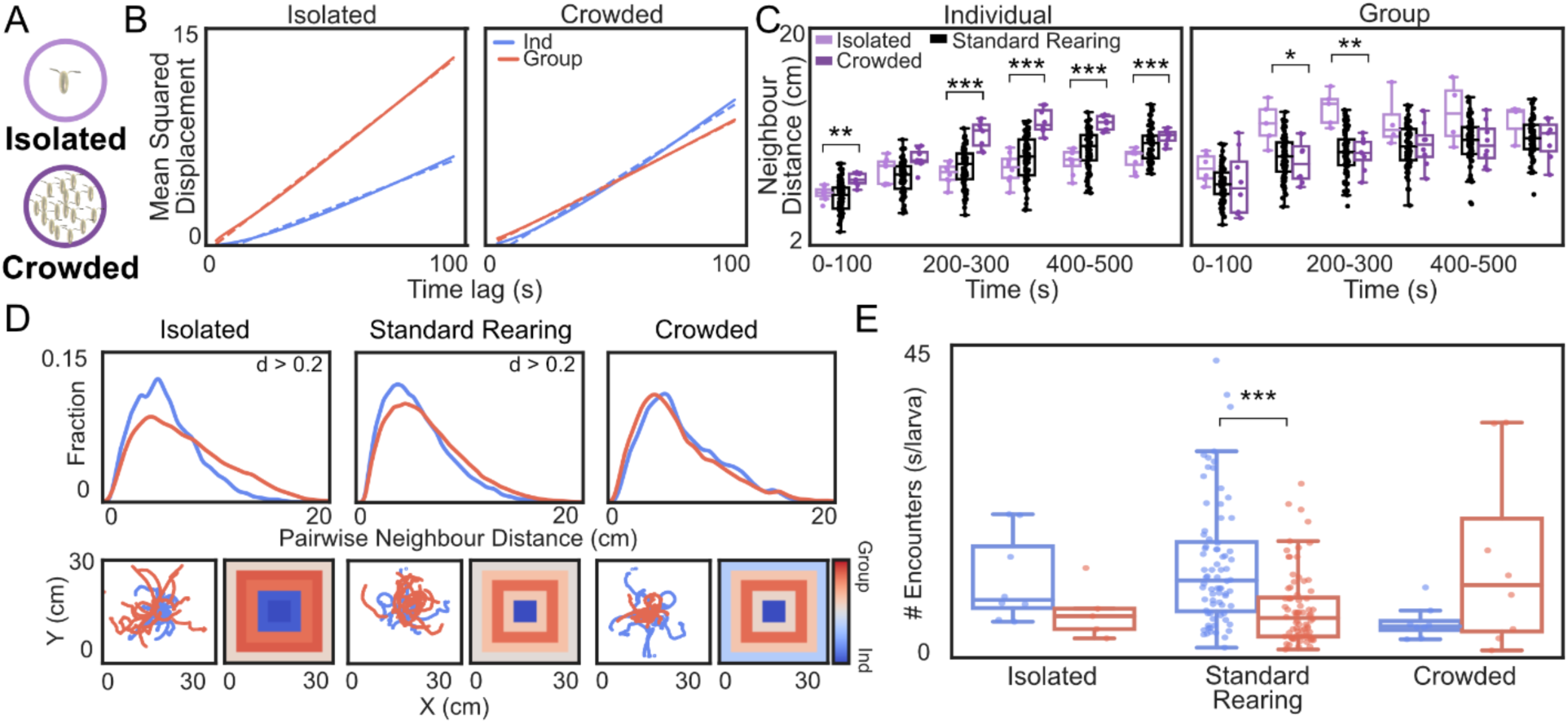
Developmental experience influences social modulation: (**A**) Larvae were grown either in isolation from a single egg (bright purple) or in crowded conditions (dark purple, 50 eggs per well). (**B**) Mean squared displacement was plotted for each treatment for the superimposed groups (N = 8 per treatment, blue) and real groups (N = 5 for isolation; N = 8 for crowded, red). Standard Rearing = same data as in Figure 1. (**C**) Average neighbor distance over time for the superimposed and real groups. Statistic comparison was performed between isolated and crowded groups, which were measured in parallel. (**D**) Top: Pairwise neighbor distance for the first 100s. Effect size = Cohen’s d (0.2 < d < 0.5 – small). Bottom: Sample trajectories and distribution heatmaps. (**E**) Encounter frequency in the real and superimposed groups over the first 100s. (Mann-Whitney U test = * p < 0.05; ** p < 0.01, *** p < 0.001).

Compared to standard rearing conditions (same data as in Fig. 1), isolated larvae dispersed similarly when tested individually, however, ND and PD were higher when tested in a group (Fig. 3C/D). Crowded larvae showed normal ND and PD when tested in a group but spread out more when tested individually. Crowded larvae have more encounters with conspecifics in a real group compared to standard-reared larvae, whereas isolated larvae have fewer encounters in a real group (Fig. 3E). Thus, we find that developmental social experience affects larval behavior, isolation leading to increased conspecific avoidance, crowded/competitive conditions leading to higher acceptance of conspecific encounters.

### Social context modulation requires the detection of multimodal sensory cues

Larval behavior is modulated by social context; thus, we asked how larvae sense conspecifics. We leveraged the genetic toolkit available in *Drosophila* to test sensory receptor mutants (Fig. 4A) and larvae with ablated sensory neurons (Fig. 4B, Table S1). To ablate sensory neurons, we crossed the sensory neuron GAL4-driver lines with *UAS-hid,reaper* (*32*) to induce apoptosis (*33*). Individual larvae were tested in parallel to control for locomotion deficits or other behavioral changes (Fig. S2A/B, S3A-K). We identified that different sensory modality impairments affect social context modulation in real group experiments (Fig. 4A/B).

**Fig. 4:**
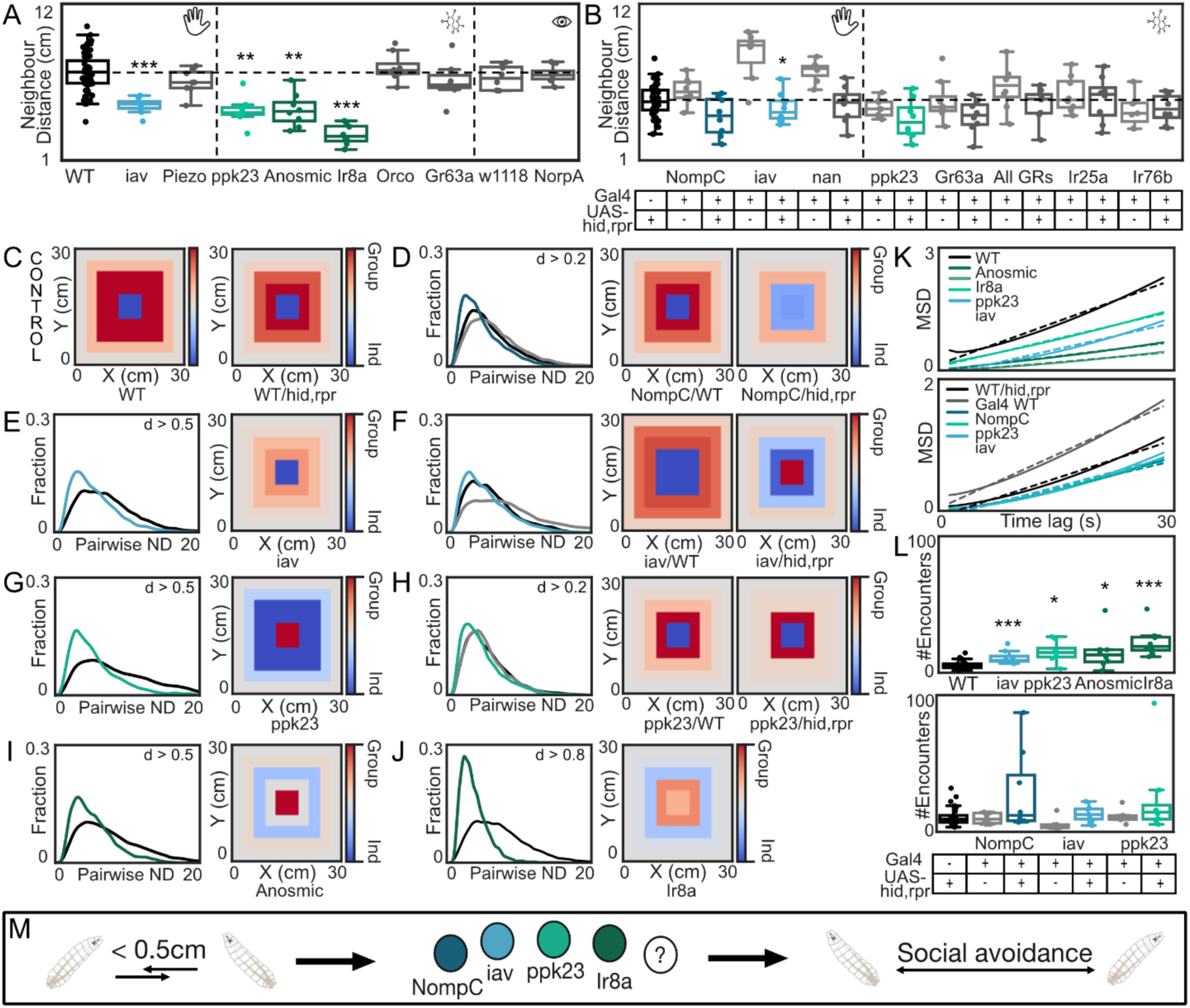
Conspecifics are recognized by multimodal sensory cues: (**A**) Mutant experiments: Real group average neighbor distance (ND) for the first 100s. WT results are pooled over all experiments (black, N = 54). Mutant lines with no phenotype (grey, N = 8) and mutant hits (different colors, N = 8). (**B**) Genetic silencing: Real group average ND for the first 100s. *WTx hid-reaper* (indicated by hid,rpr) control results are pooled over all experiments (black, N = 40). Experimental cross/GAL4 control with no phenotype (grey, N = 8). Experimental cross hits (different colors, N = 8). (**C**) Distribution heatmaps for the pooled controls: WT (N = 24) and *WT x hid/reaper* (N = 24). (**D-J**) Pairwise neighbor distance for the first 100s in all real groups. Effect size = Cohen’s d (0.2 < d < 0.5 – small; 0.5 < d < 0.8 – moderate; d > 0.8 – high). Controls = black/grey, mutants/experimental groups = different colors. Distribution heatmaps (superimposed groups (blue) and real groups (red)). (**K**) Mean squared displacement (MSD) for all hits/controls. (**L**) Encounter frequency for all hits. (**M**) Social context modulation is mediated via four different mechano– and chemosensory channels and receptors. (Mann-Whitney U test = * p < 0.05; ** p < 0.01, *** p < 0.001).

Larvae with impaired mechanosensation (inactive = iav, No mechanoreceptor potential C = NompC) show reduced ND. The mechanosensitive channels iav and NompC have been shown to detect the touch and vibration of conspecifics in adults (*15*, *23*). Knockout of the mechanosensitive piezo channel or ablation of nanchung (nan) expressing neurons did not affect social dispersal (Fig. 4A/B).

Impairing chemosensation (Pickpocket23 = ppk23, Ionotropic receptor 8a = IR8a (anosmic)) also causes larvae to have reduced ND. Anosmic mutants (*23*), which have mutations in *orco*, *Ir8a, IR25a,* and *GR63a*, show a similar reduction in the neighbor distance as *IR8a* mutants (*23*). As we find no impairment in social distancing in the *orco* mutant, *IR25a x hid,reaper,* and *GR63a* mutant (see also Fig. S2A/B, S3A-K), we conclude that the anosmic phenotype is due to the *IR8a* mutation. ppk23 is a receptor involved in sensing larval deposits and egg pheromones (*27*, *28*). We find that it is also required for normal social dispersal behavior in fly larvae. The w1118 control, blind *NorpA* mutants (*34*), and silencing of all gustatory receptors (all GRs) (*35*) or IR76b expressing neurons (*36*) showed no difference to WT behavior in real groups.

Real group NDs for the WT control and the *WT x hid,reaper* control did not differ across experiments, thus Fig 4A/B includes a pooled dataset (Fig. S2A/B). We also visualized the average positional distribution of the WT control and the *WT x hid-reaper* control in a single heatmap plot (Fig. 4C). All hits have a low PD and spread out less than individuals (Fig. 4D-J). The ppk23 receptor mutant phenotype was stronger than the genetic silencing experiment, this could be explained by an incomplete GAL4 expression pattern or weak GAL4 expression by the driver line. Also, in all hits, larvae in real groups have a lower dispersal rate (Fig. 4K) and a higher encounter frequency in the first 100s, indicating that they stay closer together (Fig. 4L). This is however not due to locomotion defects, as individually tested larvae show no difference in pairwise neighbor distance nor dispersal in the arena (Fig. S2C-J). In summary, we find that larvae recognize conspecifics through multimodal sensory cues via touch, ppk23, and Ir8a receptors (Fig. 4M).

## Discussion

We find that conspecific exposure affects *Drosophila* larval behavior in a non-food context. Larvae explore the arena differently when they are in a group compared to when they are alone. The effect of social context on larval behavior is robust and prioritized over other external cues, such as aversive light and attractive sugar. Isolated larvae show an enhanced dispersal in the presence of conspecifics; thus, larvae adapt to their social environment during development. Finally, we show that conspecifics provide multimodal sensory cues and are recognized through mechanosensory and chemosensory pathways.

### Dispersal in the group context

In a non-food context, larvae usually explore an experimental arena as they search for food. We show that individual larvae show slow exploration and perform local search patterns – short runs and bigger turns (Fig. S1A). However, when tested in a group, larvae disperse faster and perform straight, fast runs away from conspecifics after an encounter. Encounters with alive conspecifics seem required to induce social modulation, as adding obstacles, such as agar pieces, into the arena does not enhance dispersal in individual larvae (Movie S4, Fig. S1B) (*26*). Most conspecific encounters take place in the first 100s of the experiment (Fig. 1E), however, larval dispersal behavior is modulated over the whole 10-minute experiment (MSD, Fig. 1C), thus our data suggest that the initial exposure to conspecifics induces a shift in the internal state that lasts over many minutes. Social modulation might be beneficial in the absence of food, as larvae compete for potential food sources and might themselves become a food source for conspecifics (*22*). Indeed, in the presence of protein-rich and familiar fly food, larval dispersal is reduced and we find no social modulation (Movie S3, Fig. S1A). This is in line with previous studies, which show that groups of larvae even perform collective digging in normal fly food (*19*).

### Social cues vs. external cues

We tested individuals and groups of larvae in different sensory contexts and found that individual larvae respond stronger to an attractive fructose substrate or an aversive light source (Fig. 2). Individual larvae dispersed less on fructose, probably due to immediate exploitation of the sweet and nutritious fructose substrate. Grouped larvae dispersed similarly on both substrates, thus, social context overwrote the response to fructose. Fructose is sweet and attractive, however, it is not an optimal food source for larvae as they require protein for development (*37*). Individual larvae dispersed faster when we illuminated the arena with white LED lights. We hypothesize that this is due to the stress caused by aversive light (*31*). As the LED lights were placed at the arena borders, they created a weak light gradient towards the middle of the arena. Therefore, the individual larvae returned to the middle of the arena as they were avoiding the illuminated borders. However, grouped larvae dispersed similarly in the illuminated or dark arena, thus, social context also overwrote the response to light over the whole time of the experiment. These results show that social context can override responses to other sensory inputs and is a strong modulator of larval behavior. We however also find that stronger external stimuli, such as protein-rich food (Video S3, Fig. S1), override the social modulation effect of the conspecifics. This might also be possible with other highly salient stimuli, such as pure odors (*38*, *39*) or ON/OFF light choices (*31*). Future experiments will reveal how social context is integrated with other sensory cues and how it can suppress certain behavioral responses.

The social modulation effects are highly reproducible in our big arena (25cm x 25cm). Many other behavioral larval group assays use a smaller arena size (⌀ 10cm), restricting the exploration range. Thus, even though the social modulation effect might be less obvious when testing groups or individuals in these smaller assays, conspecifics might still influence larval responses to weak external cues.

### Social experience

We tested whether social context modulation is an innate behavior or whether it is shaped during development. Testing isolated larvae that never have encountered another larva before in a group revealed an increased social modulation effect (Fig. 3). Thus, during normal development, larvae often encounter conspecifics in the presence of food and might thus be less aroused by conspecific exposure. Flies can learn about the conspecific presence and show decreased stress responses afterward (*24*); the same might be true for larvae. In our crowded condition, grouped larvae behaved similarly to our standard-reared group; however, when isolated, they dispersed faster. This might be due to competition, stress, or reduced food access in crowded conditions. Investigating how food deprivation affects individual and grouped larval behavior will require further examination.

Generally, animals exhibit plasticity in their sensory processing and behavioral decisions in response to isolation or conspecifics (*40–44*). Thus, uncovering how neural circuits adapt to social experiences in *Drosophila* larvae might also reveal general principles and mechanisms that can be adapted to other species.

### Conspecifics are recognized via multimodal sensory cues

Our results suggest that encounters with conspecifics seem to be the main driving force of social modulation and the induction of a social state. Therefore, we looked at how larvae sense conspecific presence. We find that mechanosensory (NompC, iav) and chemosensory (ppk23, IR8a) receptors are required for social modulation. The mechanosensory neurons expressing NompC and iav have been shown to be required for sensing gentle touch and vibrations in adult flies and larvae (*15*, *19*). Thus, fly larvae are probably capable of sensing the touch or vibrations of other live larvae. ppk23 is known to be required for larval deposit preference and mediates repulsion to fly eggs, preventing egg cannibalism (*27*, *28*). In adult flies, ppk23 is involved in courtship behavior (*45*). We find that ppk23 is necessary for social dispersal in larvae, supporting its role in mediating aversion in an alive larva context (*28*). We hypothesize that ppk23 may generally be required for species recognition but in a context-dependent manner. It may drive aversion in the presence of live conspecifics when no food is available (*28*) while promoting attraction when food or attractive cues are present but no other alive conspecifics (*27*). Finally, we observe a strong phenotype in IR8a/anosmic mutants. While previous literature suggests no known IR8a expression in larvae, sensitive PCR experiments indicate the presence of *IR8a* mRNA (*36*, *46*). IR8a is required for acid sensing in adult flies and mosquitoes (*47–50*). Further investigation is needed to elucidate the potential role of Ir8a in mediating social interactions in larvae.

We cannot exclude that additional receptors or sensors might be involved in conspecific recognition; we might not have found a phenotype in our screening due to redundant requirements. In future experiments, silencing multiple receptors might reveal such additional sensors. In summary, larval conspecifics are recognized via multimodal sensory pathways, similar to what was found in adult flies (*15*). Uncovering the sensory systems required for social context-dependent modulation paves the way for dissecting the central brain circuits involved in conspecific recognition and understanding how social cues are integrated with and can overwrite other sensory processing pathways.

## Conclusion

Animals make complex behavioral decisions based on their internal state (*51*), experience (*24*), sensory context (*40*, *52*, *53*), and social context (*23*). We show that conspecifics also affect larval *Drosophila* behavior in a non-food context and the presence of environmental cues. These findings highlight the general importance of social exposure during development and accounting for social influence in group experiments.

Social cues are important in our daily lives and social isolation can severely impact human health (7) *Drosophila* shares homologous genes and conserved neural subtypes with all animals. The available genetic toolkit and the whole-brain connectome in *Drosophila* offer a distinct advantage over other taxa as they enable precise dissection of the neural circuits underlying social interactions. This will facilitate the investigation of the neural basis of social behavior and health in other taxa ranging from insects to mammals.

## Materials and Methods

### Animal stocks and husbandry

Flies were raised on a standard cornmeal diet and kept in incubators at 25^0^C and 60% humidity on a 12-hour light/dark cycle. Adult flies were allowed to lay eggs for 48 hours and were then removed from the vials. The eggs were allowed to develop for 4-6 days. Larvae in the middle to late second instar stage were used for experimentation.

### Transgenic lines

Mutant fly lines and crosses for experiments were maintained as mentioned above and crossed with a specific number of animals to maintain similar crowding during development. For a single cross, 10 GAL4-driver line males and 25 UAS-hid/reaper female virgins were collected and allowed to mate. Control crosses for both the UAS-hid/reaper and the GAL4-driver line crossed to WT were also made with a similar sex ratio. Wildtype experiments were performed with the Canton-S strain. Stocks were obtained from Bloomington *Drosophila* Stock Center or courtesy of other labs (BDSC, see Table S1).

### Behavioral Experiments

#### Social modulation

Either a single larva or a group of 15 larvae were allowed to roam freely on a 25 cm x 25 cm assay plate filled with 100 ml of a 2% agarose substrate. The arenas were maintained at a constant temperature of 25^0^C and 60% humidity and placed in a light-tight box. Larval behavior was recorded for 10 minutes with a Basler camera (acA2040-90umNIR) and lens (Kowa Lens LM16HC F1.4 f15mm1”) at 1fps with a red light filter (Edmund optics #89-837) above the arena. The arena was illuminated with red light using infrared LEDs from the bottom (Solarox® LED strip infrared 940 nm).

For all behavioral experiments, middle to late second-stage instar larvae were used of a similar size (4-6 days after egg laying). Larvae were removed from the fly vials and washed in distilled water to remove all traces of food. To study individual behavior, a single larva was placed in the middle of the assay plate. These experiments were repeated with different larvae at least 30 times. To study group behavior, 15 larvae, if not stated otherwise, were placed in the middle of the agarose plate. These experiments were repeated at least 8 times, if not stated otherwise.

When testing larval behavior in the presence of fly food, a small piece of food (1-2g of standard cornmeal food) was placed in the center of the arena. 15 larvae were placed in the center and this was repeated 8 times.

To test if physical encounters are sufficient to enhance dispersal, we placed physical non-larval obstacles, 14 pieces of 2% agar (3-4mm) in the center of the arena. An individual larva was placed in the center and the experiment was repeated 30 times.

#### Change in arena context

To test if an attractive substrate influences social modulation, larvae were tested on a substrate containing fructose. Therefore, we mixed agarose with D-(-)-Fructose (Merck, CAS #57-48-7) and poured it into the square arena plates (for one plate: 36g/100ml = 2M).

To test whether the light environment influences social modulation, larvae were tested in an illuminated arena. Therefore, we placed strips of white light LEDs (Solarox® 12V LED strip reel warm white) on three sides of the experimental box, which illuminated the agarose plate from the side. One side was not illuminated, as this was the side where the arena was closed with a black curtain. No specific side preference was detected in the larvae.

#### Isolation assay

Flies were allowed to lay eggs for 24 hours on standard fly food in a fly cage. The eggs were collected the next day and placed in separate wells containing standard fly food in a 48-well plate. For the isolation treatment, a single egg was placed in one well. Each well was closed with a cotton plug to prevent larval interactions. The food and cotton plug were sprayed with water to prevent the drying out of the food. For the crowded treatment, 50 eggs were placed in a single well. The survivorship in this procedure was only around 50%, probably because of the limited availability of food and space. All plates were maintained at a constant temperature of 25^0^C and humidity of 65%. After 4-6 days, the larvae were removed from the well plates and tested using the assay setup as described before.

### Quantification and Statistical Analysis

#### Behavioral Analysis

Videos were analyzed using the freely available tracking software TRex (*29*) to obtain the position (X/Y), speed, and midline offset of individual larvae. To quantify larval dispersal, we calculated the mean squared displacement (MSD) for specific time intervals. The slope of the MSD curve provides insights into the diffusion properties of the larval groups.

To visualize the position of the larvae, we plotted their location in a probability heatmap. The heatmap was made by subtracting the position data of individually tested larvae from the real group position data. After subtracting, the values of each region were averaged out in concentric squares to focus on the distance from the center, the initial position of the larva. A positive value indicates the higher probability of finding a larva from a group at this location (red), and a negative value indicates the more likely presence of individuals (blue). Based on X/Y coordinates, the average neighbor distance of each larva was calculated to all its neighbors in the group. For individual larvae, an artificial group was formed by randomly selecting 15 larvae, and the neighbor distance was calculated as described above. The pairwise neighbor distance was also calculated between each pair of larvae and plotted as a kernel density estimate plot to visualize the distribution of the distances between the pairs.

To quantify local interactions, we counted larval encounters, which were defined by a larva pair with less than 0.5cm distance from each other. Encounter frequency was calculated as the number of frames in which a larva encounters another larva. Encounter duration was calculated by adding consecutive frames within a single encounter (seconds/encounter). The speed and bending angle of larvae were calculated during and after (5s) the encounter. Box plots or density plots are shown to visualize the data. Box plots indicate the first and third quartiles with the median. Whiskers indicate the 5 and 95 percentiles of the data. Dots indicate group averages from individual experiments.

#### Statistical Analysis

To detect significant differences between experimental groups (ND, Encounter frequency), we performed a non-parametric Mann-Whitney U test, which can be employed even for not normally distributed data. The test was conducted with treatment (social context, environment or genotype) as the grouping factor. To compare pairwise neighbor distance across groups, we calculated the Cohen’s d effect sizes to quantify the magnitude of differences between the two groups. The values of effect size were interpreted as high (d > 0.8), moderate (0.5 < d < 0.8), small (0.2 < d < 0.5) and negligible (d < 0.2).

## Acknowledgments

The authors would like to thank Andreas Thum, Iain Couzin, Einat Couzin-Fuchs, and Armin Bahl for valuable discussions. We further thank Andreas Thum and Simon Sprecher for providing fly lines.

## Author contributions

Conceptualization: AM, KV. Methodology: AM, KV. Investigation: AM, EZD. Visualization: AM. Supervision: KV. Writing—original draft: AM, KV. Writing—review & editing: AM, EZD, KV.

## Data and materials availability

All data needed to evaluate the conclusions in the paper are present in the paper and/or the Supplementary Materials. Additional data related to this paper will be available on request.

## Supplementary Information

**Table S1.** Fly lines used in this study. *Kindly provided by the Thum lab

| Genotype | Source |
| --- | --- |
| UAS-hid, reaper | * (32) |
| iav<sup>-/-</sup> | * BDSC #24768 |
| Piezo [KO] | * BDSC #58770 |
| Δppk23 | * BDSC #33300 |
| Anosmic | (23) |
| Orco [1] | BDSC #23129 |
| Gr63a [1] | * BDSC #9941 |
| Ir8a [1] | * BDSC #41731 |
| NorpA | * BDSC #9047 |
| w1118 | * BDSC #3605 |
| NompC-Gal4 | BDSC #36361 |
| iav-Gal4 | * BDSC #52273 |
| nan-Gal4 | * BDSC #24903 |
| ppk23-Gal4 | * BDSC #93026 |
| Gr63a-Gal4 | * BDSC #9942 |
| All GRs-Gal4 | Kindly provided by the Sprecher lab<br>(GO-Gal4) (35) |
| Ir25a-Gal4 | BDSC #41728 |
| Ir76b-Gal4 | * BDSC #41730 |

**Fig. S1:**
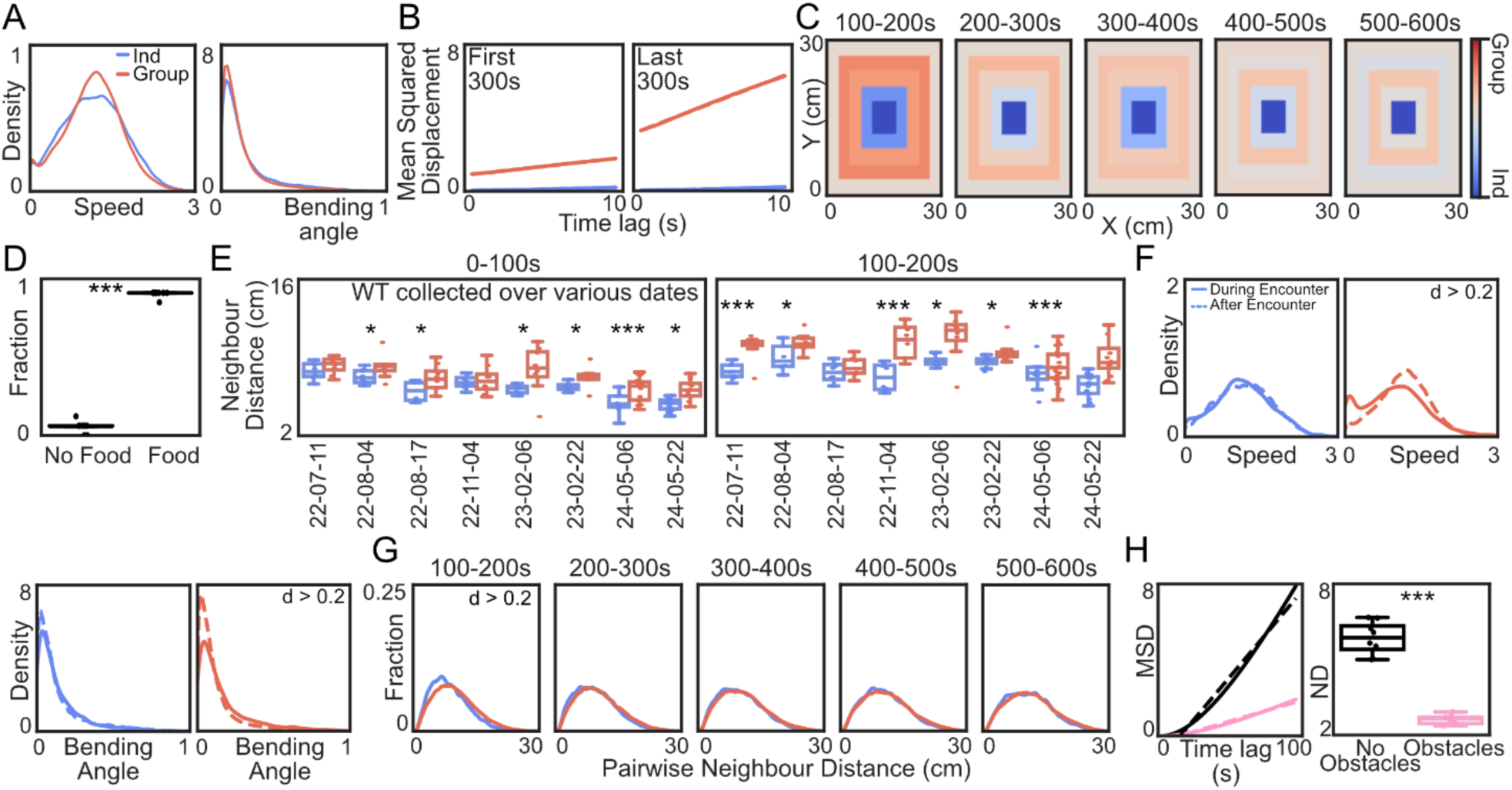
Behavioral analysis for larvae as individuals and in a group: (**A**) Speed and bending angle for superimposed groups (blue) and real groups (red). (**B**) Mean Squared Displacement (MSD) for real and superimposed groups for the first 300s and last 300s of the experiment. (**C**) Heatmaps for WT data from 100-600s of the experiment in 100s intervals. (**D**) Fraction of larvae in the center with food or no food placed in the center of the arena (N = 8 trials). (**E**) The average neighbor distance of superimposed individual groups (blue) and real groups (red) for WT data collected over various time points. (Mann-Whitney U test = * p < 0.05; ** p < 0.01, *** p < 0.001). (**F**) Speed and bending angle for superimposed groups (blue) and real groups (red) during (line) and after encounter (dashed line). (effect size = Cohen’s d (d < 0.2 – negligible, 0.2 < d < 0.5 – small, 0.5 < d < 0.8 – moderate, d > 0.8 – high)) (**G**) Pairwise neighbor distance for WT data from 100-600s of the experiment in 100s intervals. (**H**) MSD for individual larvae in the absence (black)and presence (pink) of agar obstacles (N = 30 trials). Average neighbor distance (ND) of superimposed individual larvae (bootstrapped to N=8 trials) for both treatments.

**Fig. S2:**
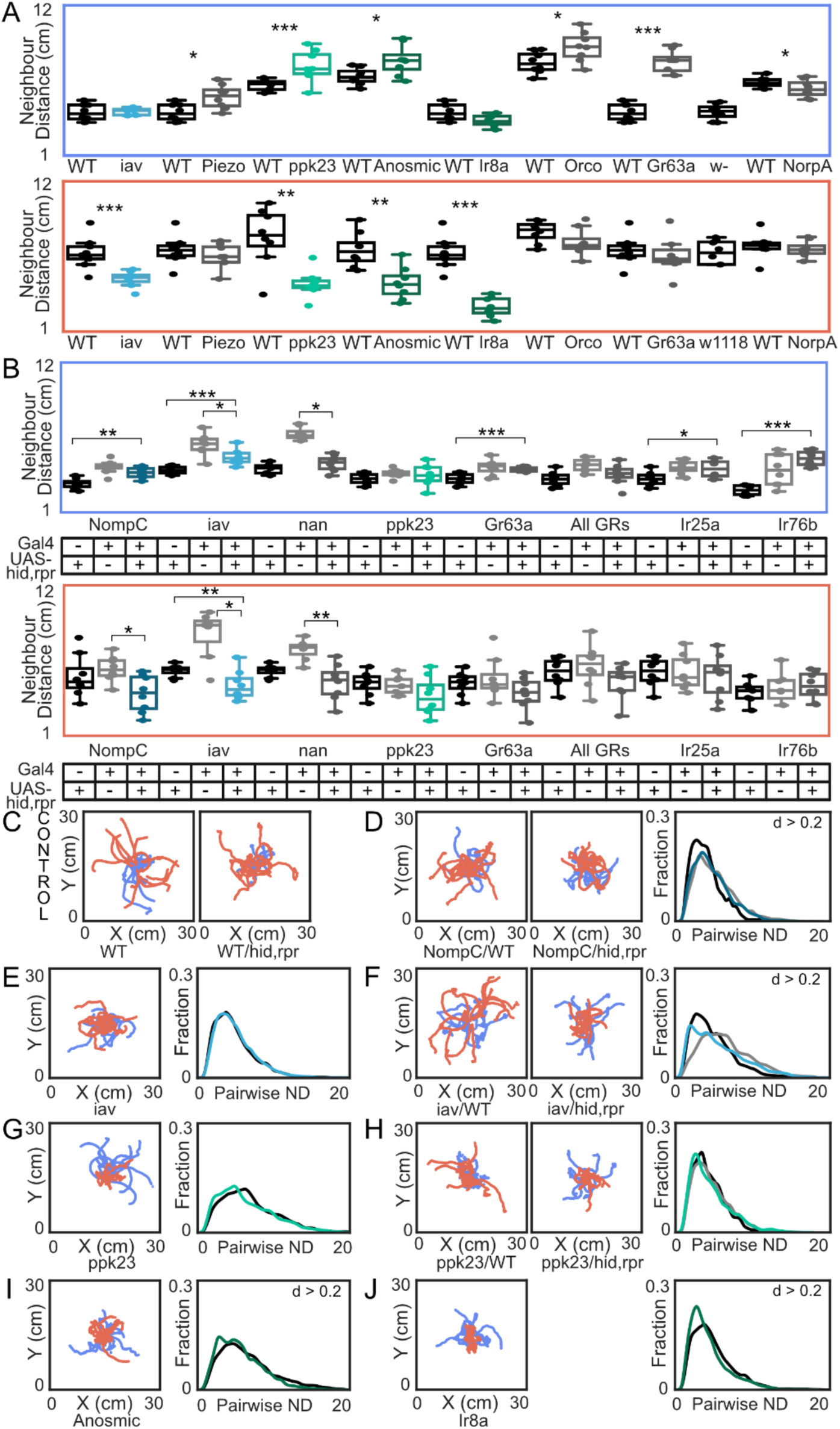
Summary of all genetic screenings and behavioral analysis for the superimposed individual groups with social phenotype: (**A**) Mutant experiments: Superimposed individual groups (blue box) and real groups (red box) for the first 100s. WT control (black, N = 8) and mutant lines with no phenotype (grey, N=8) and with phenotype (different colors, N = 8). (**B**) Genetic silencing: Superimposed individual groups (blue box) and real groups (red box) for the first 100s. *WT x hid,reaper* (black, N = 8), experimental controls/GAL4 controls with no phenotype (grey, N = 8), and cross hits (different colors, N = 8). (**C**) Sample trajectories of WT and *WT x hid,reaper* for a superimposed individual group (blue) and real group (red). (**D-J**) Sample trajectories. Pairwise neighbor distance for the first 100s in all superimposed individual groups. Effect size = Cohen’s d (0.2 < d < 0.5 – small).

**Fig. S3:**
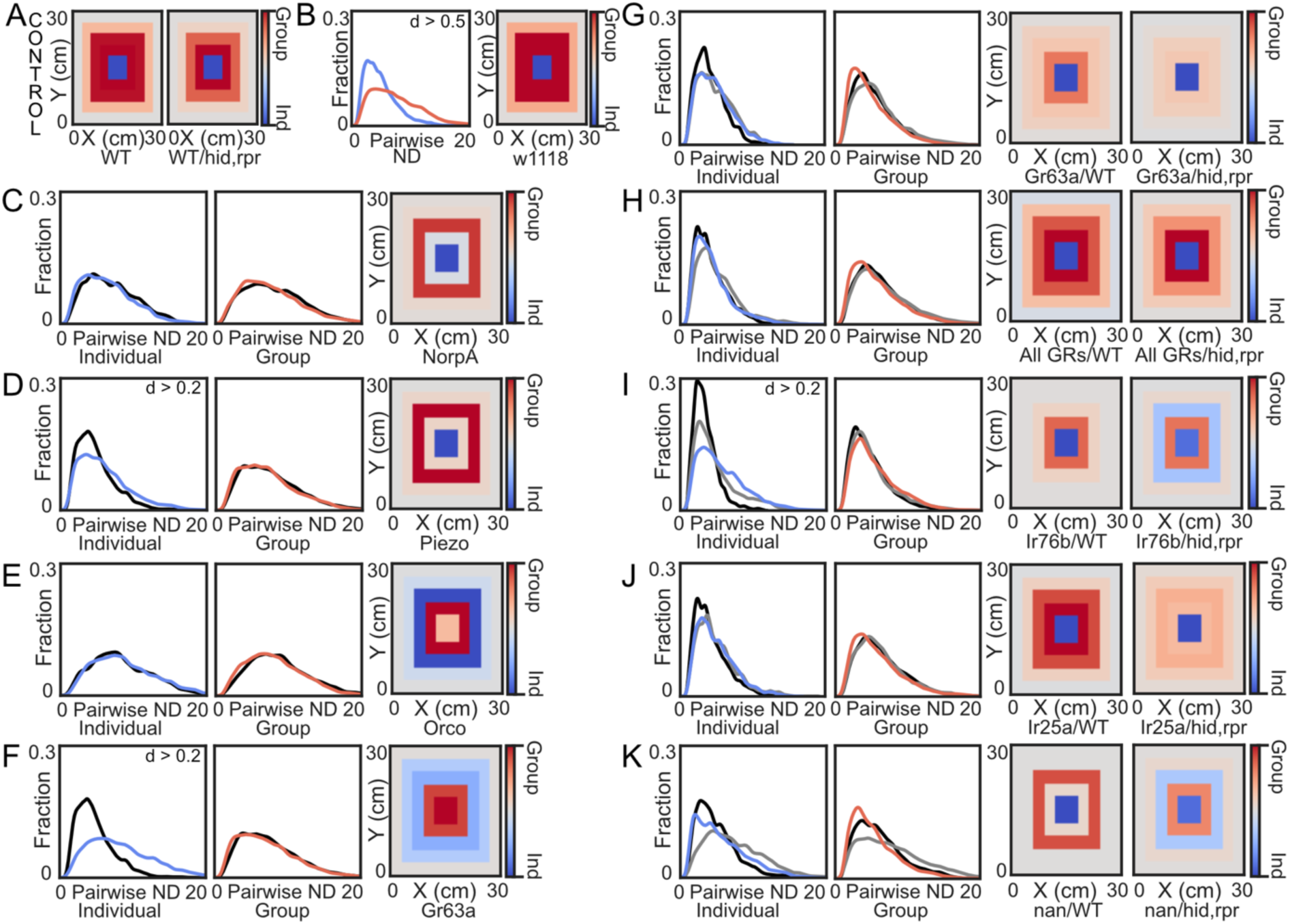
Behavioral data for all the genetic screenings with no phenotype: (**A**) Distribution heatmaps for the pooled controls: WT (N = 32) and *WT x hid,reaper* (N = 32). (**B**) Pairwise neighbor distance for the first 100s of superimposed individual groups and real groups for w1118 larvae (N = 8). Effect size = Cohen’s d (0.2 < d < 0.5 – small, 0.5 < d < 0.8 – moderate). Distribution heatmap for w1118 larvae. (**C-K**) Pairwise neighbor distance for first 100s in all superimposed individual groups and real groups. Controls = black/gray, Experimental groups = blue (superimposed individual group) / red (real group). Distribution heatmaps for all controls and treatment groups.

## Supplementary Videos

**Movie S1**.

Dispersal behavior of larvae in a group of 15 on agar for the first 100s.

**Movie S2**.

Dispersal behavior of an individual larva on agar for the first 100s.

**Movie S3**.

Dispersal behavior of larvae in a group of 15 exposed to standard cornmeal food for the first 100s.

**Movie S4**.

Dispersal behavior of individual larvae exposed to agar obstacles for the first 100s.

